# Validation of ligand tetramers for the detection of antigen-specific lymphocytes

**DOI:** 10.1101/2022.10.24.513230

**Authors:** Kristin S. Fitzpatrick, Hanna N. Degefu, Katrina Poljakov, Madeleine G. Bibby, Allison J. Remington, Tyler G. Searles, Matthew D. Gray, Jim Boonyaratanakornkit, Pamela C. Rosato, Justin J. Taylor

**Author notes:** Contact, @JustinTaylorLab.

## Abstract

The study of antigen-specific lymphocytes has been a key advancement in immunology the past few decades. The development of multimerized probes containing antigens, peptide:MHC complexes, or other ligands was one innovation allowing the direct study antigen-specific lymphocytes by flow cytometry. While these types of studies are now common and performed by thousands of laboratories, quality control and assessment of probe quality is often minimal. In fact, many of these types of probes are made in-house and protocols vary between labs. While peptide:MHC multimers can often be obtained from commercial sources or core facilities, few such services exist for antigen multimers. To ensure high quality and consistency with ligand probes, we have developed an easy and robust multiplexed approach using commercially available beads able to bind antibodies specific for the ligand of interest. Using this assay, we have sensitively assessed the performance of peptide:MHC and antigen tetramers and find considerable batch-to-batch variability in performance and stability over time. This assay can also reveal common production errors such as miscalculation of antigen concentration. Unexpectedly, probes including the fluorochrome allophycocyanin exhibited more rapid performance decline compared to probes including the fluorochrome R-phycoerythrin when both were stored at 4°C. This performance decline was reduced for most, but not all, batches when antigen tetramers were instead stored at −20°C in 50% glycerol. This work could set the stage for the development of standardized assays for all commonly used ligand probes to limit lab-to-lab technical variation, and experimental failure due to probe underperformance.

## INTRODUCTION

Over the past several decades, the use of fluorochrome-conjugated ligands of all types has become a standard approach to detect antigen-specific lymphocytes by flow cytometry. For antigen-specific B cell detection, the most used tools are fluorochrome-conjugated antigen/peptide tetramers, fluorochrome-conjugated virus-like particles as well as other types of fluorochrome-conjugated ligands (For an incomplete cross-section, see (1–232)). Combined, the use of these types of tools has led to important advances in all fields of immunology, the development of experimentally and/or clinically useful antibodies, and as a starting point for rationally designed vaccine antigens.

Throughout the course of a study analyzing antigen-specific B cells by flow cytometry, it is often necessary to produce dozens of antigen probe batches to conduct experiments over the long periods of time research often takes. Even with strict production protocols, this often means that different individuals have produced different batches of antigens using different lots of reagents, or even different reagents all together. Antigens can also undergo storage for different amounts of time at different temperatures during the various production steps, or multiple freeze/thaw cycles as protocols often neglect to set strict guidelines for every factor that could influence antigen structure. These issues are magnified across fields, where the same antigen could be produced, purified, and stored using completely different protocols. For example, one researcher may produce an antigen using a bacterial system while another used a mammalian system, which could influence glycosylation, folding, or stability of the antigen. Alternatively, production and purification of an antigen may in one case include multiple freeze/thaw cycles while another protocol would not, which could alter the stability of the antigen probe. Another difference can occur after probe production, when one researcher may produce an antigen tetramer fresh for each experiment, whereas a different researcher stores the tetramer for different amounts of time at 4°C or −20°C before use. Despite these sources of variation, there are typically few validation steps to ensure consistent and reproducible antigen probe performance within each experiment. Here, we describe a rapid and robust approach to validate the quality of B cell antigen and MHC tetramers before and/or within each experiment that can be used to authenticate this key resource.

## RESULTS

### Comparison of murine B cells binding different batches of an antigen tetramer

As an initial assessment, we tested whether different batches of SARS-CoV-2 RBD APC tetramers would bind the same frequency of murine B cells. These different batches were made using the same protocol and either used the same day or stored for up to 42 weeks before use. For this, spleen and lymph nodes were pooled from multiple animals and divided into three technical replicates for each tetramer batch to gauge technical variability (Fig. 1a). Cells were next incubated with SARS-CoV-2 RBD APC tetramers and a control APC-DyLight755 tetramer that allows for cells binding streptavidin-APC to be excluded from the analysis. Following incubation, tetramer-binding B cells were enriched from the sample using anti-APC microbeads to allow for more robust visualization. Using this approach, ~5% of B cells in the APC-enriched fractions bound to SARS-CoV-2 RBD APC tetramer batches made the day of the experiments, which amounted to ~0.08% of B cells in samples without enrichment (Fig. 1b, c). Likewise, a similar frequency of cells can be found using some SARS-CoV-2 RBD APC tetramers produced as many as 42 weeks earlier when stored at −20°C in 50% glycerol (Fig. 1c, d). Tetramers stored at 4°C in 1xDPBS performed poorly after only a few weeks, as demonstrated by over 30-fold fewer cells binding to tetramers from these batches. Importantly however, storage at −20°C in 50% glycerol did not maintain consistent cell binding of tetramer for all batches (Fig. 1c, d). Analysis of batch 7G revealed that 0.06% of B cells bound this tetramer 6 weeks after production, which was at the lower end of the range found with fresh tetramers (Fig. 1c, d). However, this frequency declined to 0.015% 17 weeks after production, and 0.0031% of cells 29 weeks after production (Fig. 1d). Batch 1G also exhibited a decline in cell binding over time, albeit at a rate slower than that of 7G (Fig. 1d). Together, these data indicate that while storage conditions can help to maintain tetramer performance over time, assessment of these tools is still important to ensure performance.

**Figure 1.**
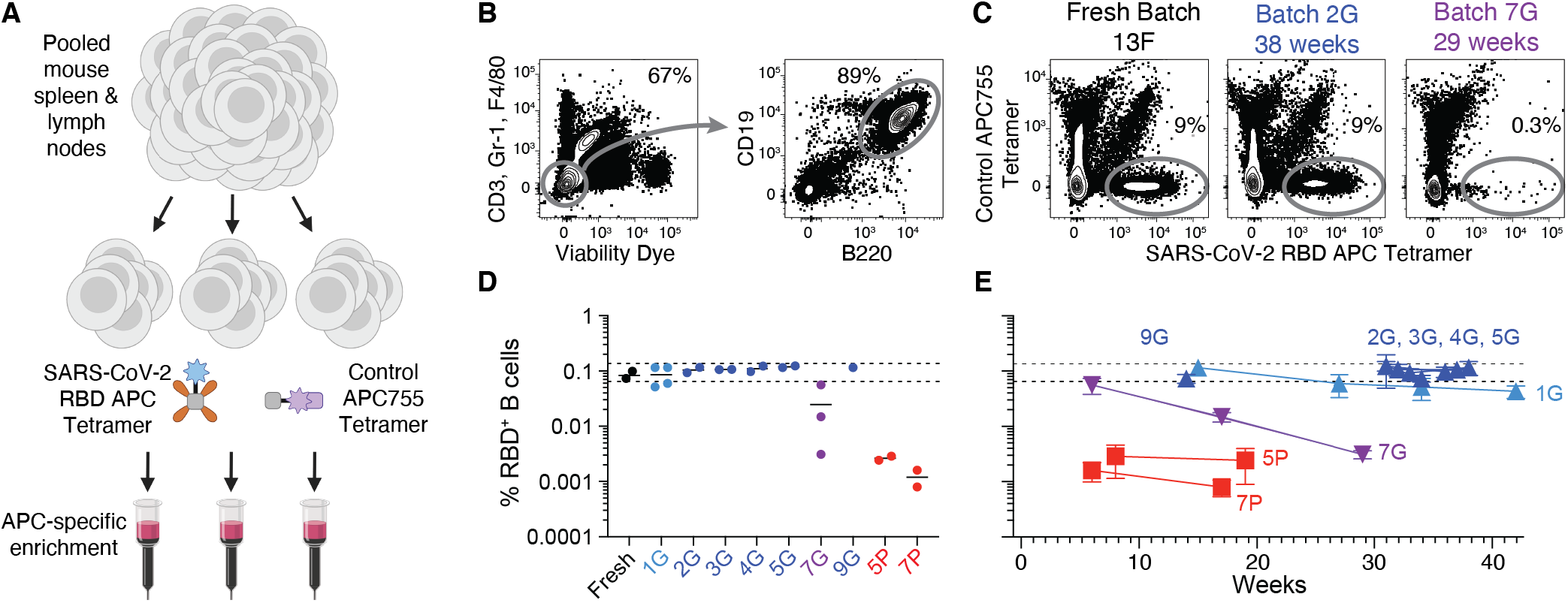
Assessing the frequency of antigen-specific B cells detected using different batches of antigen tetramers. (**A**) Schematic representation of experiments to measure the frequency of antigen-specific murine B cells bound by different batches of SARS-CoV-2 RBD APC tetramers. Spleen and lymph nodes from two to four C57BL/6 mice were pooled and divided evenly into three technical replicates for each tetramer assessed. The samples were next incubated with RBD APC tetramers and a control APC-DyLight755 (APC755) tetramer, followed by magnetic enrichment of tetramer-binding B cells using APC-specific microbeads. The control APC755 tetramers are included to gate out B cells specific for streptavidin-APC. (**B**) Representative flow cytometric gating of live CD19^+^ B220^+^ CD3^-^ Gr-1^-^ F4/80^-^ B cells from an APC-enriched fraction. (**C**) Representative flow cytometric gating of B cells binding to three different batches of RBD APC tetramers, a freshly made batch called 13F, a batch made 38 weeks prior called 2G, and a batch made 29 weeks later called 7G. The % of B cells in the APC-enriched fractions is displayed on the plots. (**D**) Combined data from six experiments showing the frequency of B cells in the starting sample that bound to two freshly made batches of RBD APC tetramers, and 9 batches made previously and stored for various amounts of time at −20°C in 50% glycerol (batches ending with G), or 4°C in 1xDPBS (batches ending with P). Each data point represents the average of three technical replicates and the lines represent means. The dotted lines represent the highest and lowest frequency of RBD-binding B cells obtained using freshly made batches. (**E**) Same data displayed in **D** except displayed versus the amount of time tetramers were stored prior to each use. Data points are displayed along with standard deviation from technical replicates.

### Bead-based assay to validate tetramer performance

Given the laborious nature of using cell samples to validate tetramer performance, we aimed to develop an easy and robust assay that could be used before, or within every experiment. For this, we expanded upon previously developed bead-based assays (233; 132). For this, eComp Ultra plus compensation beads were incubated with monoclonal antibodies specific for SARS-CoV-2 RBD, positive control antibodies specific for APC, or negative control antibodies specific for PE (Fig. 2a). Following incubation with antibodies, the beads were washed and incubated with tetramer prior to analysis. Using this approach, we found that RBD APC tetramer batch 2G was bound strongly by both anti-APC and the anti-SARS-CoV-2 RBD clone S309 (234), but not anti-PE (Fig. 2b). In contrast, tetramer batch 2P was not bound by S309 but maintained binding by anti-APC (Fig. 2b).

**Figure 2.**
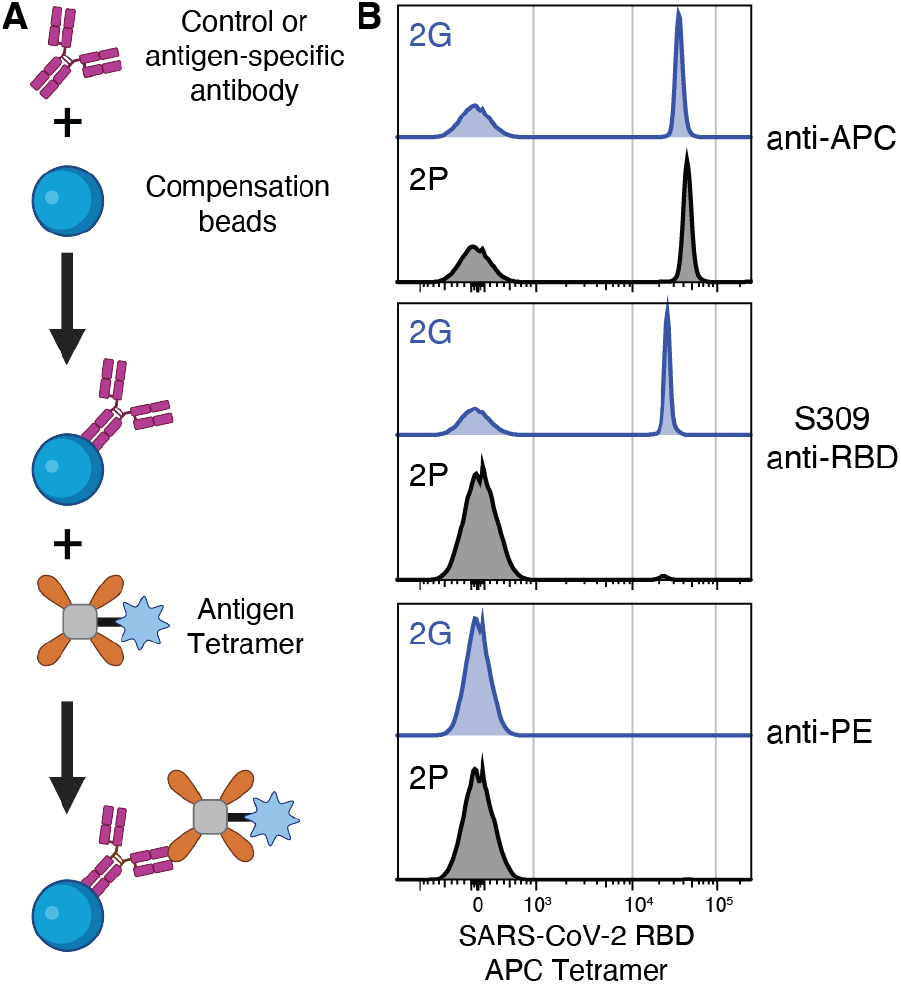
Bead-based approach to assess antigen tetramer performance. (**A**) Schematic representation of an assay in which compensation beads are loaded with S309 SARS-CoV-2 RBD-specific antibodies, positive control APC-specific antibodies, or negative control anti-PE antibodies prior to incubation with RBD APC tetramers. (**B**) Representative flow cytometric analysis of two different batches of RBD APC tetramers. These batches were produced as a single batch that was split and stored at −20°C in 50% glycerol (2G) or 4°C in 1xDPBS (2P) 36 weeks prior to analysis.

### Identification of additional antigen-specific monoclonal antibodies

We next reasoned that assessing tetramer binding by a single antigen-specific antibody clone was not enough to fully validate tetramer performance and expanded the analysis to include additional antibody clones with varied epitope specificities and binding characteristics. For this we used a combination of RBD-specific antibodies identified by others previously (S309, CV30 (235), 204, 208, 211, 215 (182)) as well as antibodies we identified specific for a SARS-CoV-2 spike antigen. For the latter, APC tetramers containing a stabilized spike trimer called S2P (236) were used to identify SARS-CoV-2 spike-specific B cells in APC-enriched PBMCs from convalescent individuals (Fig. 3a). From this screen, six S2P-binding antibodies, named E1, E7, E11, 2B5, 2B8, and 2B11, were identified and used in this study (Fig. 3b). Five of these antibodies bound to RBD with notable differences in off-rate (Fig. 3b,c). The sixth antibody, 2B5, did not bind RBD (Fig. 3b) and was used as a negative control in subsequent RBD tetramer binding experiments. The mAbs 215 and 208 bound RBD weakly compared to the other antibodies (Fig. 3c).

**Figure 3.**
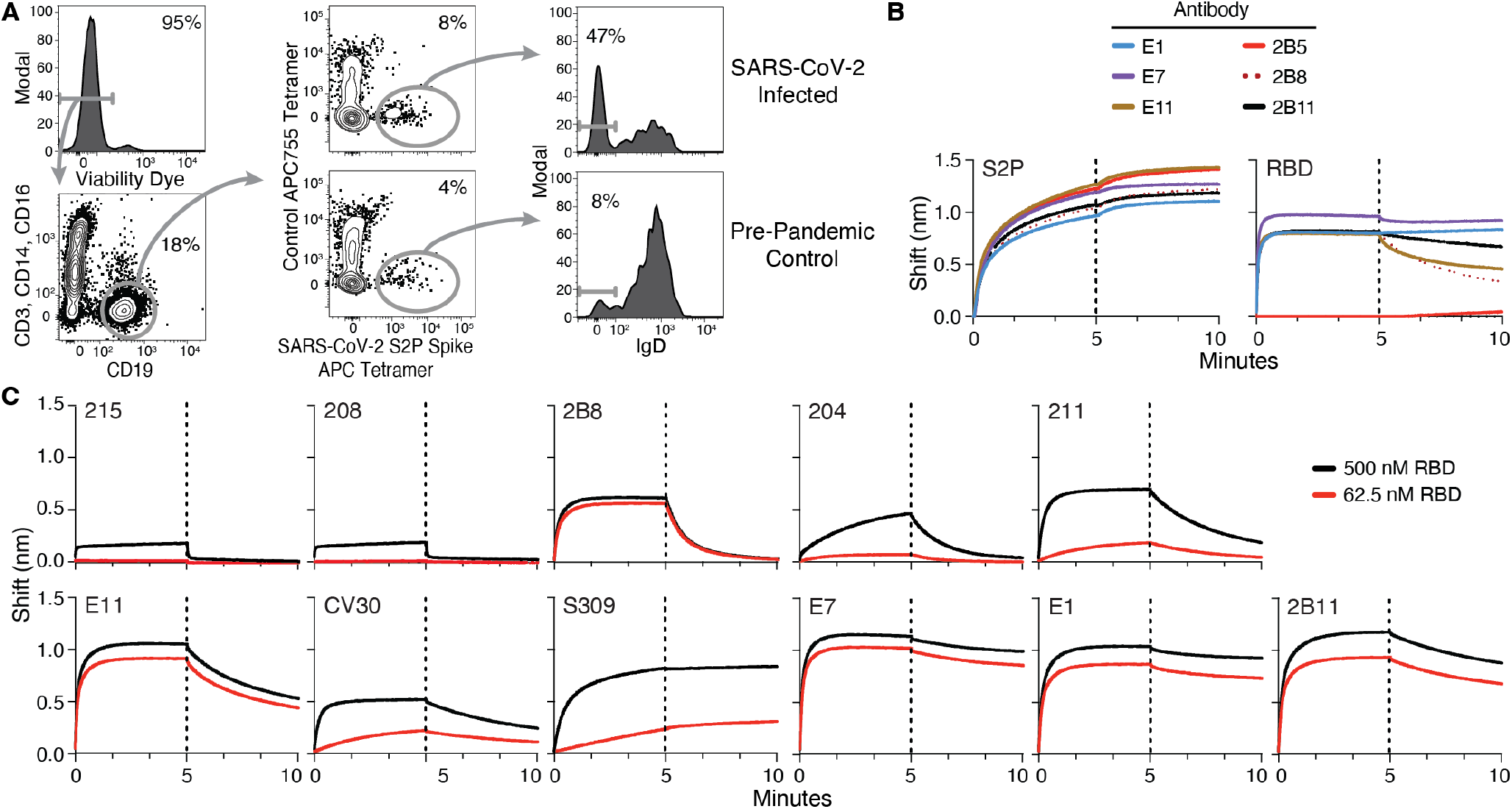
Identification and assessment of antibodies specific for SARS-CoV-2 spike. (**A**) Representative flow cytometric gating of live CD19^+^ CD3^-^ CD14^-^ CD16^-^ B cells binding SARS-CoV-2 Spike S2P APC tetramers but not APC755 tetramers from PBMCs collected from individuals hospitalized with COVID-19 three-six weeks earlier, and a control sample collected prior to the COVID-19 pandemic. Cells from three COVID-19 patients were concatenated for display. To identify spike-specific antibodies that responded to infection, single S2P-tetramer binding B cells were sorted into individual wells of a 96-well plate and underwent nested RT-PCR followed by Sanger sequencing to identify paired heavy and light chain sequences which were expressed as secreted IgG1. (**B**) Antibodies from six S2P tetramer binding B cells were assessed for binding to S2P or RBD using Bio-Layer Interferometry (BLI). (**C**) BLI assessments of binding of five RBD-binding antibodies from **B** (E1, E7, E11, 2B5, 2B8 & 2B11), and six RBD-specific antibodies from published studies (CV30, S309, 215, 208, 204 & 211) to 62.5 or 500 nM of RBD.

### Multiplexed bead-based tetramer validation assay

To assess the binding of RBD tetramers to multiple RBD-specific and control antibodies, aliquots of beads were co-incubated with an antigen-specific or control antibody as well as an irrelevant fluorochrome-conjugated antibody (Fig. 4a). The fluorochrome-conjugated antibody was included to allow the different antibody-labeled bead populations to be pooled prior to tetramer labeling and distinguished during analysis (Fig. 4b). Using this approach, we included 8 RBD-specific antibodies, anti-APC, anti-PE, and an additional negative control antibody specific for SARS-CoV-2 spike outside of the RBD. After gating on the individual bead populations, we found that different RBD-specific antibodies bound different amounts of RBD-APC tetramers as assessed by gMFI. In general, there was no correlation in the level of tetramer binding of the antibodies bound to beads (Fig. 4c) and the strength of the binding by BLI (Fig. 3c). This disconnect may be the result of avidity gains due to tetramerization of RBD in the bead assay, but not BLI assessment.

**Figure 4.**
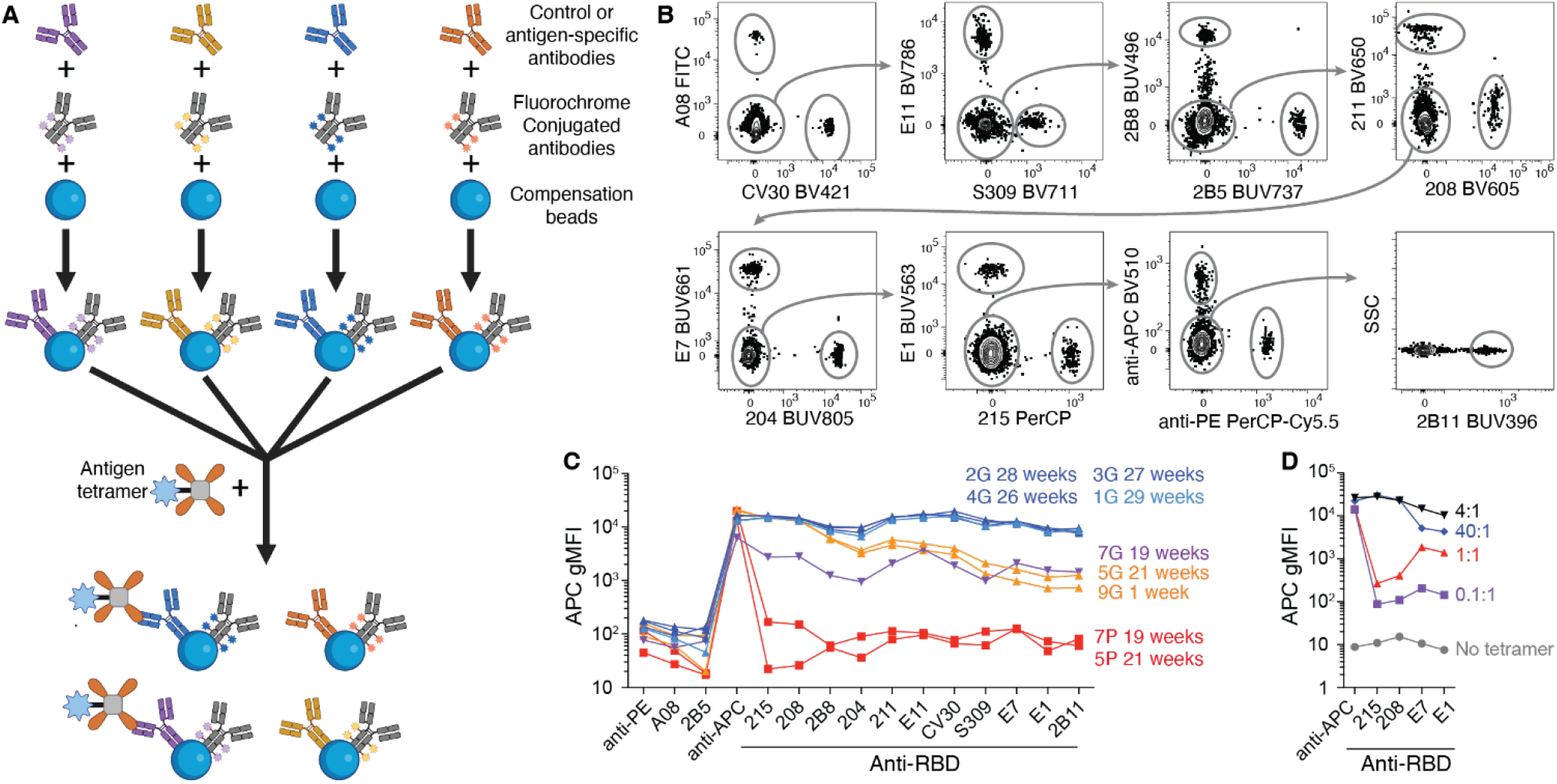
Multiplexed bead-based approach to assess antigen tetramer performance. (**A**) Schematic representation of an assay in which different populations of compensation beads are loaded with different RBD-specific or control antibodies along with unique fluorochrome-labeled antibodies prior to bead pooling and incubation with RBD APC tetramers. (**B**) Representative flow cytometric gating of fifteen bead populations in a pooled sample. RBD-specific antibodies in the panel are: 215, 208, 2B8, 204, 211, E11, CV30, S309, E7, E1, and 2B11. Negative control antibodies include 2B5, A08 and anti-PE; anti-APC is used as a positive control. (**C**) APC gMFI values of antibody-loaded beads bound to nine different batches of RBD APC tetramers stored for the listed amount of time at −20°C in 50% glycerol (batches ending with G), or 4°C in 1xDPBS (batches ending with P). (**D**) APC gMFI values of antibody-loaded beads bound to batches of antigen probes made by mixing 40, 4, 1, or 0.1 pmole of RBD-biotin to 1 pmole of streptavidin-APC.

When comparing binding in this assay to multiple batches of tetramers, four distinct profiles emerged. The first profile included most of the tetramer batches that were bound by ~0.08% of murine B cells in Figure 1. These tetramers, batches 1G-4G, were bound well by all RBD-specific antibodies and anti-APC (Fig. 4c). The second profile included batches 5P and 7P, which were the tetramer batches bound poorly by murine B cells in Figure 1. Both 5P and 7P exhibited ~100-fold lower binding by all RBD-specific antibodies (Fig. 4c), consistent with poor performance in cell binding assays in Figure 1. The third profile was displayed by batch 7G, which exhibited ~10-fold reduction binding to all RBD-specific antibodies 19 weeks after production (Fig. 4c), which is after this batch began performing poorly in enrichment experiments (Fig. 1e).

The final profile was shown by batches 9G and 5G, which exhibited poor binding by some of the RBD-specific mAbs but was bound well by others (Fig. 4c). Interestingly, no reduction in binding was detected for 215 or 208, and only slight reductions in binding for 2B8 and 204 (Fig. 4c). These antibodies also exhibited the poorest binding to RBD when assessed by BLI (Fig. 3c), suggesting that lower affinity antibodies bound batches 9G and 5G well while higher affinity antibodies could not. Upon assessment of the production of batch 9G, it was discovered that the concentration of antigen used to produce this tetramer was 10-fold higher than the 4:1 antigen:streptavidin ratio normally used to produce tetramers. Since the tetramers used in these experiments were not purified prior to use, this means that there were ~36 excess unbound antigen molecules per tetramer molecule contaminating these preparations. Therefore, it seems likely that this unlabeled free antigen was bound by high affinity antibodies, interfering with tetramer binding and detection. Lower affinity antibodies would be expected to be less affected by the presence of free antigen since stable binding is not easily achieved without multimerization. While we did not discover any production errors for batch 5G, we suspect that a similar issue occurred during preparation. Interestingly, batches 5G and 9G were assessed in cellular experiments, and bound ~0.08% of cells, similar to tetramer batches that were performing well (Fig. 1). To confirm that excess free antigen could block binding of high affinity antibodies but not low affinity antibodies, we produced batches in which a 40:1 ratio of RBD-Biotin to streptavidin-APC was compared to a 4:1 ratio. In these experiments, beads loaded with the high affinity antibodies E1 or E7 exhibited reduced binding to the 40:1 batch compared to the 4:1 batch (Fig. 4d). In contrast, beads loaded with the low affinity antibodies 208 or 215 bound similarly to the 40:1 batch and 4:1 batch (Fig. 4d). Together, these results indicate that excess free antigen in tetramer preparations can block the binding of high affinity antibodies, but not binding of lower affinity antibodies expressed by majority of cells in repertoires without no previous SARS-CoV-2 RBD exposure.

In these experiments we also explored how probes containing fewer than 4 antigen molecules would perform compared to their fully loaded tetramer counterparts. For this we compared probe batches made with RBD-biotin:Streptavidin ratios of 0.1:1 and 1:1, compared to properly made 4:1 RBD tetramers. As expected, the beads loaded with the low affinity antibodies 208 and 215 exhibited ~100-fold loss in binding to the 1:1 batch, which was further reduced at 0.1:1 (Fig. 4d). In contrast, binding by beads loaded with the high affinity antibodies E1 and E7 to the 1:1 batch was only ~10-fold decreased (Fig. 4d). Together, these data highlight the utility of using a bead-based assay to reveal errors in antigen tetramer production resulting in underloaded probes.

### Assessment of antigen tetramer stability over time

Using a cocktail of beads bound to different antibodies, we conducted experiments assessing the performance of different batches of tetramers over time under different storage conditions. In total, four batches of RBD APC or RBD PE tetramers were produced, split, and stored at −20°C in 50% glycerol, or stored at 4°C in 1xDPBS. Assessing the APC tetramers revealed that all batches stored at −20°C in 50% glycerol largely maintained stable binding by RBD-specific antibodies over time (Fig. 5). In contrast, RBD APC tetramer batches stored at 4°C in 1xDPBS began to exhibit reduced binding by RBD-specific antibodies as early as 1-3 weeks after production when assessed with CV30, S309 and 2B11 (Fig. 5b). By six weeks after production, most batches stored at 4°C had 5-10-fold reduced binding (Fig. 5b). Antibodies such as 2B8 and E11 also exhibited a decline in binding with slightly slower kinetics (Fig. 5b). These results highlight the importance of using multiple antibodies to assess tetramer performance. Unexpectedly, the batch-to-batch decline in performance varied widely, with batch 2 only exhibiting a moderate loss in binding when stored at 4°C for the 12 weeks assessed. Importantly, the decline in performance was not detected equivalently by each of the RBD-specific antibodies in this experiment.

**Figure 5.**
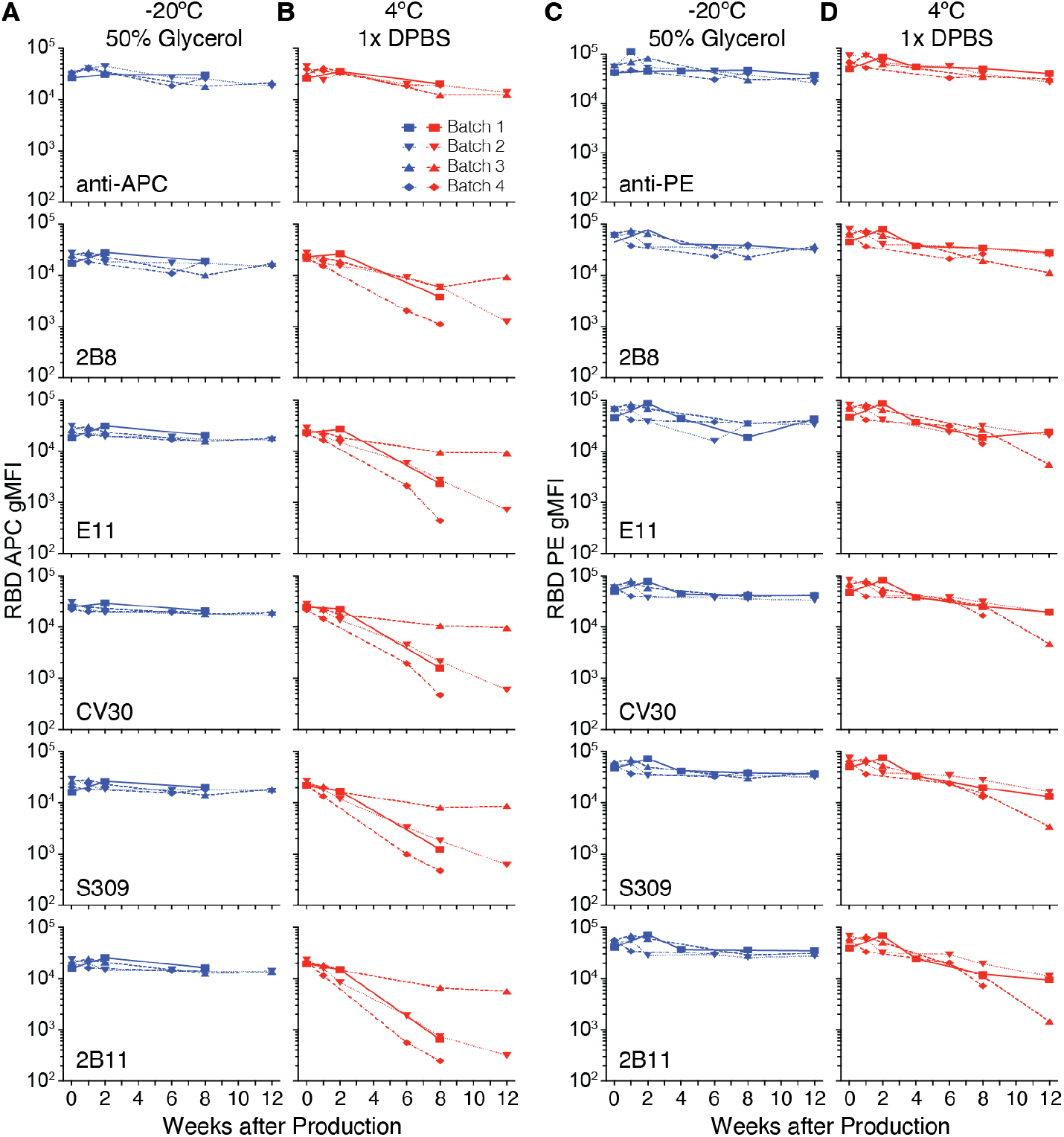
Assessing antigen tetramer performance over time. Combined data from 10 experiments in which four batches of RBD APC tetramers (**A**, **B**) and RBD PE tetramers (**C**, **D**) were assessed using a multiplexed bead assay containing beads loaded with antibodies specific for RBD (2B8, E11, CV30, S309, 2B11) and anti-APC. Each batch of RBD APC and RBD PE tetramers was split in half and stored at −20°C in 50% glycerol (**A**, **C**) or 4°C in 1xDPBS (**B**, **D**).

### Antigen tetramer stability with different fluorochromes

We initially predicted that the decline in tetramer performance over time when stored at 4°C in 1xDPBS reflected the stability and half-life of the antigen under these conditions. However, this did not appear to be the case, since RBD PE tetramers were bound well by anti-RBD antibodies for weeks longer than RBD APC tetramers made at the same time (Fig. 5c-d). Overall, these data suggest that PE-conjugated tetramers are more stable at 4°C compared to APC-conjugated tetramers.

### Adapting the assay for a wider array of fluorophores

Since APC and PE are not the only fluorochromes used in experiments using ligand tetramer-binding cells, we adapted this assay to include additional positive controls. Since antibodies specific for every possible fluorochrome are not available, we instead produced positive control beads bound to streptavidin-specific antibodies. Compared to APC-specific beads, streptavidin-specific beads bound lower levels of wild-type and omicron RBD APC tetramers (Fig. 6a, b), but at a level sufficient to serve as a positive control for omicron RBD BV421 and BV650 tetramers (Fig. 6c, d). As an additional control in these experiments, we also included beads specific to the 6x-HIS tag included in the RBDs. HIS-specific beads exhibited reduced binding to both wild-type and omicron RBD APC tetramers compared to streptavidin-specific beads, but binding was clearly detectable above the level of binding by the negative control PE-specific beads.

**Figure 6.**
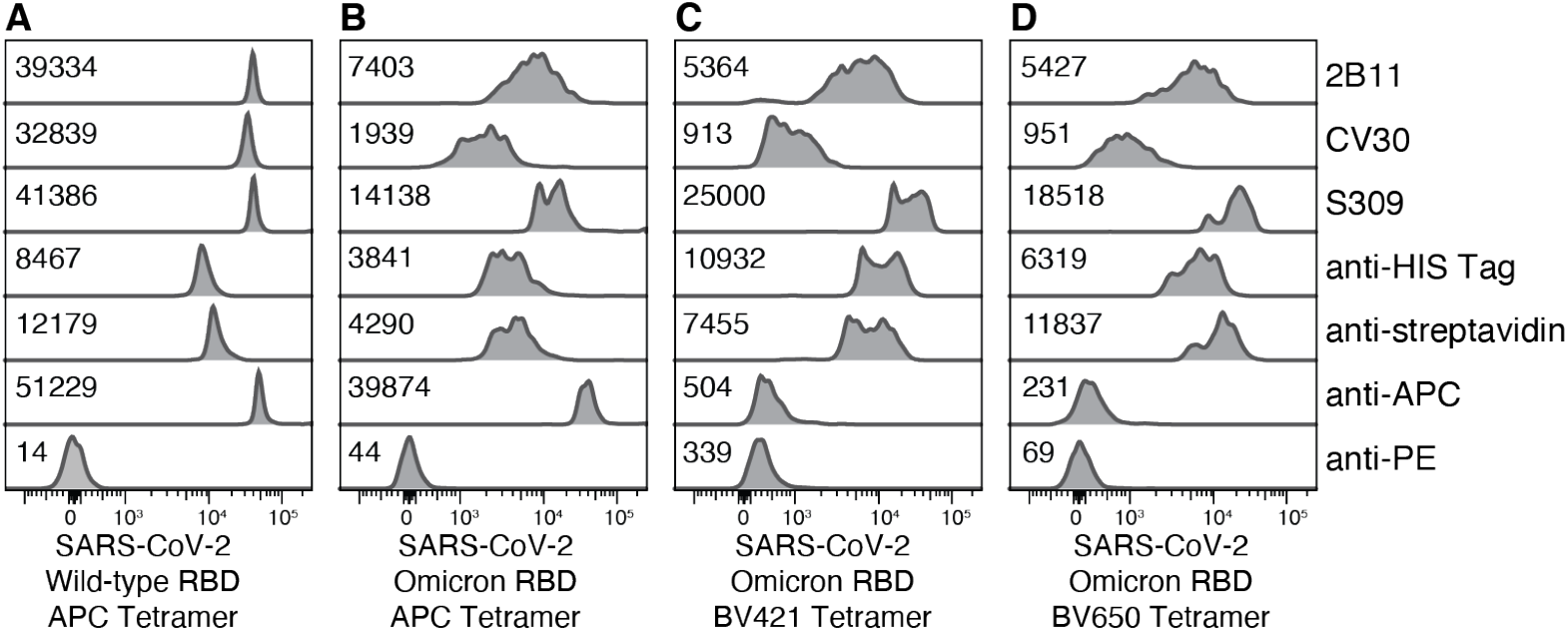
Assessing antigen tetramers utilizing other fluorochromes. Representative data from two experiments in which wild-type RBD APC tetramers, omicron RBD APC tetramers, omicron RBD BV421 tetramers, and omicron RBD BV650 tetramers were assessed for binding to multiplexed beads loaded with anti-RBD (2B11, CV30, S309), anti-HIS tag, anti-streptavidin, anti-APC, or anti-PE. The numbers on the plots represent the gMFI of APC, BV421 or BV650.

### Adapting the assay to test peptide:MHC tetramer stability

The generation of fluorescent multimerized peptide:MHC molecules (237) was a major advance in the detection of antigen-specific T cells. Many peptide:MHC tetramers are commercially available, but these tools are also commonly made by research labs. From both sources these tools can degrade over time to varying degrees. Testing tetramer integrity can be difficult, particularly in the case of human HLA tetramers where clinical samples with a reliable target T cell population may be limiting such as in the case of non-immunodominant epitopes, or cancer neoantigens. Given this, we next assessed whether our multiplexed bead-based assay could be used to test peptide:MHC tetramer stability. For peptide:MHC class I monomers to retain stability, they must include three components: β2m, a heavy chain molecule, and an 8-10 amino acid peptide. A bead-based assay using MHC-specific antibodies has been used to assess tetramer integrity, however this relied on non-commercially available beads and has not been adapted to assess multiple components of the peptide:MHC complex in one reaction (233). To test if our multiplex bead-based assay could be adapted for MHC tetramer, we used an HLA-A*02 PE tetramer loaded with the YLQPRTFLL peptide from SARS-CoV-2 spike (SARSCov2_YLQ_) (238). SARSCov2_YLQ_:HLA-A*02 monomers were made in-house by constructing HLA-A*02 monomer folded with a UV labile peptide as previously described (239–241). UV labile peptide was exchanged for the desired peptide under UV light, then the resulting monomer was tetramerized with streptavidin-PE. We incubated UltraComp beads with antibodies specific for components of the peptide:MHC tetramer complex including heavy chain (HLA-A2 or pan HLA-ABC) human β2m, anti-PE as a positive control and anti-FITC as a negative control (Fig. 7a). As described for antigen tetramers, to enable multiplexing of these bead-antibody conjugates, we included irrelevant fluorochrome-conjugated antibodies to allow the different antibody labeled-bead populations to be pooled prior to SARSCov2_YLQ_ PE tetramer labeling and distinguished during analysis (Fig. 7a). After gating on individual bead populations, we observed high PE gMFI on beads specific for PE, β2m, and HLA-A2 but not negative control beads specific for FITC (Fig. 7b&c). To confirm this assay can also be used for mouse MHC I tetramers, we incubated H-2K^b^ APC tetramer loaded with the B8R peptide from vaccinia virus (H2K^b^-B8R) (242) with bead-antibody conjugates loaded with antibodies against murine MHC I heavy chain H-2K^b^, anti-streptavidin or anti-APC as positive controls, and anti-FITC as a negative control. Consistent with results from the human SARSCov2_YLQ_:HLA-A2 test, we observed strong binding of H2K^b^-B8R APC tetramer to H-2K^b^-specific beads, streptavidin-specific beads, and APC-specific beads, but not FITC-specific beads (Fig. 7d). Together, indicating this assay can be adapted for the assessment of human and mouse MHC tetramers.

**Figure 7.**
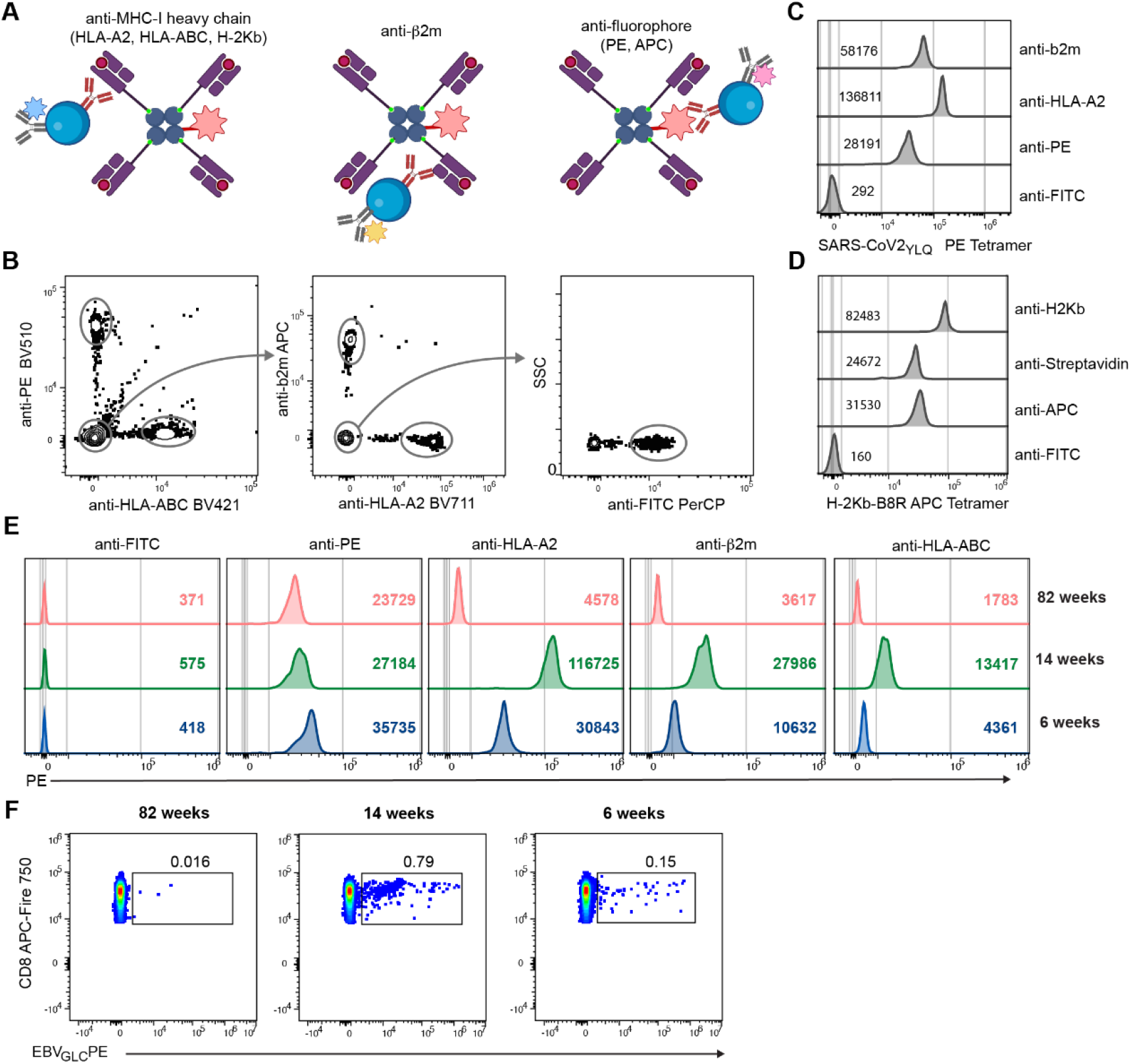
Assessing peptide:MHC tetramers using beads and cells. (**A**) Schematic representation of peptide:MHC class I tetramer binding to beads loaded with different MHC-specific or fluorophore-specific antibodies along with unique fluorochrome-labeled antibodies to distinguish bead different populations. (**B**) Representative flow cytometric gating of bead populations in pooled sample. (**C, D, E**) Flow cytometry histogram plots showing multiplexed bead staining for human SARSCoV2_YLQ_ PE tetramer (**C**) mouse B8R APC tetramer (**D**) and comparison of three different batches of EBV_GLC_ PE tetramer made 82, 14 and 6 weeks before (E). The numbers on the plots represent gMFI of PE (**C**, **E**) or APC (**D**). (**F**) Flow plots displaying the binding of EBV_GLC_ PE tetramer batches made 82, 14 and 6 weeks before by gated CD3^+^ CD8^+^ cells from the same sample of human PBMCs. Data in **B**-**F** representative of two similar experiments.

To assess batch-to-batch variability and stability over time, we used the multiplexed-bead assay to compare different batches of HLA-A*02 PE tetramer loaded with the immunodominant peptide GLCTLVAML from Epstein-Barr virus (EBV_GLC_). These different tetramer batches were made using the same batch of UV labile peptide:MHC monomer, but UV peptide exchange and tetramerization were performed 82, 14 or 6 weeks earlier. The results showed a clear difference in the binding ability of tetramer made 82 weeks prior compared to more recently made tetramers. Specifically, we observed a reduction in PE gMFI with anti-HLA-A2, -β2m and -HLA-ABC antibodies, indicating a decline in tetramer integrity over time (Fig. 7e). Interestingly, we observed differences in the two recently made tetramers which did not track with time post-production; Tetramer produced 14 weeks prior showed a stronger binding profile than the 6-week-old batch (Fig. 7e). This indicates that in addition to decay over time, there is likely batch-to-batch variability contributing to tetramer integrity.

To test if the difference in binding abilities observed with beads correlates with CD8+ T cell binding ability, we stained a sample of PBMCs from an EBV+ donor with the different batches of EBV_GLC_-PE tetramer. Similar to the bead-based assay, the tetramer batch made 82 weeks prior exhibited poor performance with a very low frequency of EBV-specific T cells detected. Additionally, while the 6- and 14-week-old batches showed overall improved binding in cell-based assays, the 14-week-old batch had a brighter, more distinct population of tetramer-specific T cells compared to the 6-week-old batch, supporting the observed profile in the bead-based assay (Fig. 7f). This difference may indicate an error in tetramer production with the 6-week-old batch resulting in poor binding ability. Together, these data demonstrate the feasibility of the multiplexed-bead-based assay to test the performance of peptide:MHC tetramers.

## DISCUSSION

Here, we have developed an easy and robust assay for assessing the performance of antigen tetramers with the goal of minimizing technical variability and preventing failed experiments due to unreliable and suboptimal tools. The assay utilizes commercially available beads typically used to generate single color controls for compensation bound to monoclonal antibodies specific for the antigen in the tetramer, or MHC complex components, and control antibodies specific for the fluorochrome or streptavidin in the tetramer. By co-labeling the beads with different fluorochrome-labeled antibodies, different populations of beads can be multiplexed into a single tube and each population identified during analysis. This allows for multiple antigen-specific and control antibodies to be utilized in this assay, allowing for coverage of a wide range of binding modalities, epitopes, and in the case of MHC tetramers, components of the peptide:MHC complex. Together, this assay allowed us to detect both improperly produced tetramers, as well as tetramers that performed poorly over time.

Moving forward an assay such as we have described could be standardized to help minimize lab-to-lab differences in reagent quality and ensure reproducibility in similar experiments. Large research consortiums would be well suited to standardize the antibody screening panels for commonly used antigens, as well as controls for normalization of data generated on different flow cytometers by different researchers. Reagent supply companies could also aid this effort by producing antibody-binding beads preloaded with fluorochromes and/or pre-made cocktails targeting commonly used antigens or peptide:MHC components as opposed to the versions we produced in house.

One limitation of the described assay is that it relies upon the availability of previously characterized monoclonal antibodies specific for the antigen of interest. For newly developed or understudied antigens, these may not be readily available. However, given that antibody cloning is often an early step for tetramer validation, we would encourage researchers to utilize these cloned antibodies to assess future tetramer batches to ensure consistency with early studies. Alternatively, it may also be possible to assess tetramer performance using polyclonal antigen-specific antibodies purified from serum.

For MHC tetramer validation, we expanded on a published bead-based assay previously used to assess tetramer performance, but which has not been widely adapted (233). Our use of commercially available beads and multiplexing provides a fast and cost-effective method which may facilitate more widespread adoption of such assays for MHC tetramer validation in the future. While we focused here on MHC I tetramers, the same approaches could be adapted to test MHC II tetramer integrity using commercially available antibodies against human (HLA-DQ, -DR, and -DP) and mouse MHC II (I-A and I-E). While not tested here, antibodies specific for specific peptide:MHC combinations, which can be generated by isolating B cells specific for peptide:MHC (89), may prove even more reliable in testing peptide:MHC tetramer integrity and would enable further multiplexing to test many different tetramers in one reaction.

## MATERIALS AND METHODS

### Animals

Six- to 14-week-old C57BL/6 male and female mice were purchased from the Jackson Laboratory (Bar Harbor, ME) and maintained in a specific pathogen-free facility in accordance with Fred Hutchinson Cancer Center Institutional Animal Care and Use Committee approval and National Institutes of Health guidelines.

### Antigen Tetramers

Biotinylated wild-type RBD and biotinylated omicron RBD was purchased from Sinobiological, or produced and purified in house using the following protocol. The sequence encoding wild-type RBD was cloned into a pTT3 expression vector flanked by an N-terminal TPA signal sequence for protein secretion and a C-terminal 6x histidine-AviTag for purification and subsequent *in vitro* biotin labeling. Briefly, HEK293E cells were transfected at a density of 10^6^ cells/mL using 0.5 mg/mL of pTT3-RBD-HISAVI DNA and 2 mg/mL filter sterilized pH 7.0 Polyethylenimine (PEI) MAX (40,000 MW, Polysciences). Six days later, cells and cellular debris were removed by centrifugation at 4,000 × g for 5 minutes followed by filtration through a 0.22 μm Nalgene rapid-flow filter (ThermoFisher Scientific). Clarified supernatant was incubated with His60 Ni-Superflow resin (Takara Bio) overnight at 4°C followed by extensive washing with 0.15 M NaCl, 20 mM Tris, 10mM imidazole (pH 8.0). Antigen was eluted using 250 mM imidazole, 0.3 M NaCl and 20 mM Tris (pH 8.0). Purified protein was concentrated and buffer exchanged into 1xDPBS using a 10kDa MWCO Amicon Ultra-15 centrifugal filter (Millipore Sigma), aliquoted, and frozen in liquid nitrogen and stored at −20°C.

Aliquots of RBD-HISAVI protein were biotinylated *in vitro* using the BirA500:BirA biotin-protein ligase standard reaction kit (Avidity) according to the manufacturer’s protocol. After 2 hours incubation at room temperature, unconjugated “free” biotin was removed from RBD by extensive buffer exchanges using a 10kDa MWCO Amicon Ultra-15 centrifugal filter. Concentrated biotinylated protein was collected from the filter and ~0.1 mg aliquots were stored at −20°C.

To confirm the average number of biotin molecules bound to each antigen molecule, streptavidin-PE (PJRS25, ProZyme) was mixed with antigen in increasing ratio and incubated at room temperature for 30 minutes. Mixtures were run on an SDS-PAGE gel (Bio-Rad Laboratories) and transferred to nitrocellulose prior to incubation with streptavidin-Alexa Fluor 680 (ThermoFisher Scientific) diluted in 1x DPBS containing 0.2% Tween-20 (Millipore Sigma). The antigen:streptavidin ratio at which there was a small excess of biotinylated antigen available for streptavidin-Alexa Fluor 680 to bind was the ratio used to make tetramers with streptavidin-PE, streptavidin-APC (PJ27S, ProZyme), streptavidin-BV421 (BD Biosciences), or streptavidin-BV650 (BD Biosciences) for experiments. The molar concentration of each tetramer was calculated by measuring the absorbance of APC, BV421 or BV650 using a NanoDrop Spectrophotometer (ThermoFisher Scientific). Tetramers were stored at 0.1-1 μM in 1x DPBS at 4°C or 1x DPBS containing 50% glycerol at −20°C prior to use.

Control APC755 tetramers were created by first conjugating streptavidin-APC conjugated with NHS-DyLight 755 (APC755) following the manufacturer’s instructions (ThermoFisher Scientific). On average, Streptavidin-APC755 contained 4 - 8 DyLight755 molecules per APC. Streptavidin-APC755 was subsequently mixed with biotinylated control antigen and stored at 0.1-1 μM in 1x DPBS at 4°C or in 1x DPBS containing 50% glycerol at −20°C prior to use.

### Murine RBD Tetramer-binding B cell enrichment and flow cytometry

The spleen and inguinal, axillary, brachial, cervical, mesenteric, and periaortic lymph nodes from two-four mice were pooled, shredded using forceps, and forced through 100-micron mesh twice to generate filtered single cell suspensions. Cell suspensions were then centrifugated at 300 × g for five minutes at 4°C and supernatant discarded. Pellets were resuspended in 1.2 mL ice cold FACS buffer (1x DPBS containing 1% heat-inactivated newborn calf serum) and divided into twelve 0.1 mL fractions. Each fraction received 2 μg of anti-Fc receptor antibody 2.4G2 (Bio X Cell) and 0.75 pmol of control APC755 tetramer before incubation on ice for 5 minutes. Next, 0.25 pmol RBD APC tetramer, 0.2 μg anti-CD3 BV510 (145-2C11, BD Biosciences), 0.2 μg anti-F4/80 BV510 (BM8, BioLegend), and 0.2 μg anti-Gr-1 BV510 (RB6-8C5, BD Biosciences), 0.3 μg anti-CD19 BUV737 (1D3, BD Biosciences), 0.4 μg anti-B220 BV786 or Pacific Blue (RA3-6B2, BioLegend), and 0.5 μL Fixable Viability Stain 620 (BD Biosciences) were added and samples incubated for 25 minutes on ice. After the incubation, ~15 mL of FACS buffer was added and the samples centrifuged at 300 × g for five minutes at 4°C. The supernatant was discarded, and the pellet resuspended prior to the addition of 8.33 μL of anti-APC microbeads (Miltenyi Biotec). Following a 30 minute incubation on ice, 5 mL of FACS buffer was added and the samples were passed over a magnetized LS column (Miltenyi Biotec). The tube and column were washed once with 5 mL FACS buffer and then removed from the magnetic field. Five mL of FACS buffer was forced through the column with a plunger twice to elute column-bound cells. Following 4°C centrifugation at 300 × g for five minutes, supernatants were discarded and all of the cells from the column-bound fraction and 1/40th of the column flow through were mixed with 20,000 Fluorescent AccuCheck counting beads (ThermoFisher Scientific) to calculate the number of tetramer binding B cells in the column bound fraction, and the total number of B cells in both the column bound and flow through fraction, with the latter multiplied by 40 to account for the 1/40th of the sample interrogated. Flow cytometry was performed at the Fred Hutch Flow Cytometry Core Service on a 5-laser (355nm, 405nm, 488nm, 561nm, 640nm) LSR II, LSRFortessa or FACSymphony (BD Biosciences).

### Human S2P Tetramer-binding B cell enrichment and flow cytometry

PBMCs were isolated via venipuncture and cryopreserved from adult volunteers prior to the COVID-19 pandemic as a part of the Fred Hutchinson Seattle Area Control study or from patients diagnosed with COVID-19 as a part of NCT04338360 and NCT04344977 clinical trials. Institutional Review Board approval was obtained from Fred Hutchinson Cancer Center or the University of Washington, and all participants gave written informed consent. 10^7^-10^8^ frozen PBMCs were thawed into DMEM containing 10% fetal calf serum, 100 U/ml penicillin and 100 μg/ml streptomycin (all from ThermoFisher Scientific). Suspensions were centrifugated at 300 × g for five minutes at 4°C, supernatant discarded, and the pellet resuspended in 50 μL of FACS buffer per 10^7^ cells. Control APC755 tetramers were added at a final concentration of 25 nM in the presence of 2% rat and mouse serum (ThermoFisher Scientific) and incubated at room temperature for 10 minutes. SARS-CoV-2 S2P spike APC tetramers were then added at a final concentration of 5 nM and incubated on ice for 25 minutes, followed by a 10 mL wash with ice-cold FACS buffer. Next, 5 μL of anti-APC-conjugated microbeads were added per 10^7^ cells and incubated on ice for 30 minutes, after which, 5 mL of FACS buffer was added and the samples were passed over a magnetized LS column (Miltenyi Biotec). The tube and column were washed once with 5 mL FACS buffer and then removed from the magnetic field. Five mL of FACS buffer was forced through the column with a plunger twice to elute column-bound cells which were centrifugated at 300 × g for five minutes at 4°C and supernatant discarded. Cell pellets were resuspended to 50 μL in FACS buffer containing 6.24 μg/mL anti-IgM FITC (G20–127, BD Biosciences), 2.5 μL anti-CD19 BUV395 (SJ25C1, BD Biosciences), 1 μL anti-CD3 BV711 (UCHT1, BD Biosciences), 1 μL anti-CD14 BV711 (M0P-9, BD Biosciences), 1 μL anti-CD16 BV711 (3G8, BD Biosciences), 2.5 μL anti-CD20 BUV737 (2H7, BD Biosciences), 2.5 μL anti-IgD BV605 (IA6–2, BD Biosciences), 3.125 μg/mL anti-CD27 PE/Cy7 (LG.7F9, eBioscience), and 0.5 μL Ghost Dye Violet 510 (Tonbo Biosciences) and incubated for 25 minutes on ice. After the incubation, ~4 mL of FACS buffer was added and the samples centrifuged at 300 × g for five minutes at 4°C. Supernatants were discarded and cells resuspended in FACS buffer and tetramer-binding B cells were individually sorted using a 5-laser (355nm, 405nm, 488nm, 561nm, 640nm) FACS Aria (BD Biosciences) into empty 96-well low profile PCR plates (Labcon), which were sealed with PCR microplate sealing foil (Eppendorf), and flash frozen prior to storage at −80°C.

### B cell receptor sequencing

As described previously (243), frozen PCR plates containing individually sorted B cells were placed on ice prior to the addition of 3 μL per well of reverse transcription (RT) master mix containing 50 μM random hexamers (ThermoFisher Scientific), 0.8 μL of 25 mM deoxyribonucleotide triphosphates (dNTPs; ThermoFisher Scientific), 20 U SuperScript IV RT (ThermoFisher Scientific), 20 U RNaseOUT (ThermoFisher Scientific), 2% Igepal (Sigma-Aldrich), and RNase-free water, after which plates were incubated at 50°C for 1 hour. Following RT, 2 μL of cDNA was added to 19 μL first round PCR reaction mix containing 0.5 U HotStarTaq Polymerase (Qiagen), 0.18 μM of each 3’ reverse primer, 0.27 μM of each 5’ forward primer, 0.29 mM GeneAmp dNTPs (ThermoFisher Scientific), and 1X HotStarTaq reaction buffer (Qiagen) in RNase-free water. After the first round of PCR, 2 μL of the PCR product was added to 19 μL of the second-round PCR reaction mix containing 0.5 U HotStarTaq Polymerase, 0.18 μM of each 3’ reverse primer, 0.27 μM of each 5’ forward primer, 0.29 mM dNTPs and 1X HotStarTaq reaction buffer (Qiagen) in RNase-free water. For heavy and kappa light chains, the PCR program for both first and second round was 50 cycles of 94°C for 30 seconds, 57°C for 30 seconds, and 72°C for 55 seconds, followed by 72°C for 10 minutes. For lambda light chains, the PCR program was 50 cycles of 94°C for 30 seconds, 60°C for 30 seconds, and 72°C for 55 seconds, followed by 72°C for 10 minutes.

From the second round PCR product, 4 μL was run on an agarose gel to confirm the presence of a ~500-base pair heavy chain band or a ~450-base pair light chain band. Five μL from PCR reactions with heavy and light chain amplicons was mixed with 2 μL of ExoSAP-IT (ThermoFisher Scientific) and incubated at 37°C for 15 minutes followed by 80°C for 15 minutes to hydrolyze excess primers and nucleotides. Hydrolyzed second-round PCR products were sequenced by Genewiz with the respective reverse primer used in the second round PCR, and sequences were analyzed using IMGT/V-Quest to identify V, D, and J gene segments. Paired heavy chain VDJ and light chain VJ sequences were cloned into pTT3-derived expression vectors containing the human IgG1, Igκ, or Igλ constant regions using In-Fusion cloning (Clontech) as previously described (244).

### Monoclonal antibody production

Secreted IgG was produced by co-transfecting 10^6^ 293F cells/mL with the paired heavy and light chain expression plasmids at a ratio of 1:1 in Freestyle 293 media using 1 mg/mL PEI Max (Polysciences). Transfected cells were cultured for 7 days with gentle shaking at 37°C. Supernatant was collected by centrifuging cultures at 2,500 × g for 15 minutes followed by filtration through a 0.2 μM filter. Clarified supernatants were incubated with Protein A agarose followed by washing with IgG binding buffer (both from ThermoFisher Scientific). Antibodies were eluted with IgG Elution Buffer (ThermoFisher Scientific) into a neutralization buffer containing 1 M Tris-base pH 9.0. Purified antibody was concentrated and buffer exchanged into 1xDPBS using a 50 kDa Amicon Ultra-15 centrifugal filter.

### Bio-Layer Interferometry

Bio-Layer Interferometry (BLI) assays were performed on the Octet.Red instrument (ForteBio) at room temperature with shaking at 500 rpm. For binding analyses, anti-human IgG Fc capture biosensors (ForteBio) were loaded with 10 or 40 μg/mL of purified antibody in kinetics buffer (0.01% bovine serum albumin, 0.02% Tween 20, and 0.005% NaN_3_ in 1xDPBS, pH 7.4) for 2.5 minutes. After loading, a baseline was recorded for 1 minute in kinetics buffer. The sensors were then immersed in 62.5 or 500 nM of S2P or RBD in kinetics buffer for 5 minutes to assess association followed by immersion in kinetics buffer for 5 minutes to assess dissociation. A 1:1 binding model using ForteBio data analysis software was used for curve fitting.

### Peptide:MHC Tetramers

MHC class I heavy chain (HLA-A2) and β2m recombinant proteins were generated by overexpression in *E. coli* and purification of inclusion bodies following adapted protocol from Gabroczi *et al*. (245). To generate individual peptide:MHC monomer complexes, a refolding reaction was set up in an L-arginine rich, glutathione redox refolding buffer with protease inhibitors in the presence of high concentration of the HLA-A2 restricted UV labile peptide H-KLLT(1051)ILTI-OH (Mimotopes), where 1051 denotes a 3-Amino-3 (2-nitro-phenyl)-propionic acid residue. Briefly, peptide was dissolved in a spinning refolding buffer at 4°C prior to the slow injection of β2m followed by heavy chain protein dissolved in injection buffer (3 M guanidine-HCl, 10 mM Na Acetate and 10 mM EDTA, pH 4.2). The solution was incubated at 4°C with slow stirring for 36 hours with additional heavy chain added in a similar manner at 12 and 24 hours. After incubation, precipitated debris was removed by centrifugation at 4000 × g for 5 minutes followed by filtration through 0.22 μm bottle top filters (Corning). The monomer preparation was then concentrated, and buffer exchanged into 10 mM Tris-HCl (pH 8.0) using 10 kDa MWCO Amicon Ultra-15 centrifugal filters (Millipore Sigma). Biotinylation of monomer was carried out using BirA biotin-protein ligase kit (Avidity) according to manufacturer’s protocol. Biotinylated monomer was buffer exchanged and concentrated using 10 kDa MWCO Amicon Ultra-15 centrifugal filters prior to purification using the Superdex 200 HR 10/300 NGC Chromatography System (BioRad). The purified monomer was concentrated and stored at 0.5 mg/mL with 2 mg/mL leupeptin, 1 mg/mL pepstatin, 0.5 M EDTA and 10% sodium azide at −80°C until use. Mouse MHC tetramers were made following the same protocol using mouse MHC H-2K^b^ heavy chain protein (provided by D. Masopust) and UV labile peptide H-SIINFE(1051)L-OH (Mimotopes).

To generate peptide-specific tetramers, UV exchange reaction was set up as described in Fehlings *et al*. (241). Briefly, 62.5 μM of desired peptide and 500 μg/mL of purified monomer in 50 μL 1x PBS was incubated for 15 minutes at 4°C 4-5 cm from a 365 nm UV lamp (Analytik Jena) followed by overnight incubation at 4°C. Tetramers were created by adding Streptavidin-PE (Invitrogen) or Streptavidin-APC (BioLegend) every 10 minutes while incubating in the dark at room temperature with a final ratio of monomer to Streptavidin-fluorophore conjugate of 4 to 1. Following incubation, 400 μM of d-Biotin (Invitrogen) solution with 10% sodium azide was added to saturate any unbound biotin-binding sites on streptavidin and tetramer was stored at 4°C until use. Total protein concentration of tetramer was determined using Bradford assay kit (BioRad). Peptides used were EBV_GLC_ (GLCTLVAML, Biomatik), SARS-CoV2_YLQ_ (YLQPRTFLL, Biomatik) and Vaccinia virus B8R (TSYKFESV, Biomatik).

### Peptide:MHC tetramer cell staining

Human PBMCs were obtained from patients undergoing routine care following Dartmouth Institutional Review Board approved protocols. Cryopreserved PBMCs were thawed and rested for 2 hours in RPMI containing 10% FBS, 1% L-glutamine, 1% HEPES, 1% non-essential amino acids, 1% sodium pyruvate, 0.01% DNAse and β-mercaptoethanol at 37°C. Cells were then washed with 1x PBS and stained with LIVE/DEAD Blue stain kit (ThermoFisher) for 30 minutes on ice. After splitting the sample into three, cells were centrifuged at 400 × g for 5 minutes at 4°C. The supernatant was discarded, and the cell pellet was stained with different batches of tetramer at 1:100 on ice for 30 minutes. Next, cells were centrifuged to remove supernatant and incubated for 30 minutes on ice in 50 μL FACS buffer (1x PBS with 0.2% BSA and 10% sodium azide) containing anti-CD4 PE-Cy5 (RPA-T4, BioLegend), anti-CD8 APC/Fire750 (SK1, BioLegend), and anti-CD3 AF700 (UCHT1, BioLegend). After incubation, cells were washed twice with 3 mL FACS buffer and fixed in 2% PFA before flow cytometry.

### Antibody:bead conjugation and use

For each antibody/bead conjugate, 50 μL of UltraComp eBeads Plus compensation beads (ThermoFisher Scientific) was added to 0.125 μg of control or antigen-specific antibody and incubated for 15 minutes on ice. Two mL of FACS buffer was added and beads centrifuged at 400 × g for five minutes at 4°C. Supernatant was aspirated and beads resuspended in 50 μL of FACS buffer and stored at 4°C. For use, 5 μL of beads were mixed with 45 μL of FACS buffer containing 0.25 pmol of tetramer and incubated for 25 minutes on ice. After the incubation, ~4 mL of FACS buffer was added and the samples centrifuged at 400 × g for five minutes at 4°C. The supernatant was discarded, and the beads resuspended in 0.2 mL FACS buffer for flow cytometry.

For bead cocktails, for each antibody/bead conjugate one drop of UltraComp eBeads compensation beads was added to 0.125 μg of control or antigen-specific antibodies with 0.125 μg of fluorochrome conjugated antibodies. Four mL of FACS buffer was added and beads centrifuged at 400 × g for five minutes at 4°C. Supernatant was aspirated and in some cases beads were resuspended in 50 μL of FACS buffer prior to the addition of 100 μL of 2% paraformaldehyde and incubation for 18 minutes on ice. Four mL of FACS buffer was added and beads centrifuged at 400 × g for five minutes at 4°C. Supernatant was aspirated and beads resuspended in 50 μL of FACS buffer and stored at 4°C. For use, 3 μL of each bead type were mixed and brought up to 50 μL with FACS buffer containing 0.25 pmol of antigen tetramer or 136 μg/mL of peptide:MHC tetramer and incubated for 25 minutes on ice. After the incubation, ~4 mL of FACS buffer was added and the samples centrifuged at 400 × g for five minutes at 4°C. The supernatant was discarded, and the beads resuspended in 0.2 mL FACS buffer for flow cytometry.

Commercially available control antibodies used in this study were anti-PE (PE001, BioLegend), anti-APC (APC003, BioLegend or eBioAPC-6A2, ThermoFisher Scientific), anti-streptavidin (S10D4, ThermoFisher Scientific or Santa Cruz Biotechnology), anti-FITC (FIT-22, BioLegend), and anti-6xHIS tag DyLight680 (HIS.H8, ThermoFisher Scientific). A08 was identified via single cell antibody sequencing of human PBMC B cells and did not bind RBD when expressed as a secreted IgG1. CV30 is an RBD-specific antibody characterized previously (235), and generously provided by L. Stamatatos. 204, 208, 211 and 215 are anti-RBD-specific antibodies characterized previously (182), and generously provided by M. Pepper and J. Netland. E11, 2B8, 2B11, E1, and E7 were identified via single cell antibody sequencing of human PBMC B cells binding SARS-CoV-2 S2P spike tetramers and confirmed to bind RBD when expressed as a secreted IgG1. 2B5 was identified in the same screen and did not bind RBD when expressed as a secreted IgG1. MHC-specific antibodies used in this study were anti-human β2m (A17082A, BioLegend), anti-human HLA-A2 (BB7.2, BioLegend), anti-human HLA-ABC (W6/32, BioLegend), and anti-mouse H-2K^b^ (Y3, BioXCell).

Fluorochrome-labeled antibodies used for bead multiplexing included anti-human CD19 BUV395 (SJ25C1, BD Biosciences), anti-human CD19 BUV496 (SJ25C1, BD Biosciences), anti-human CD19 BUV563 (SJ25C1, BD Biosciences), anti-human CD79b BUV661 (HM79b, BD Biosciences), anti-human CD20 BUV737 (2H7, BD Biosciences), anti-human CD20 BUV805 (2H7, BD Biosciences), anti-human CD19 BV421 (HIB19, BD Biosciences), anti-human CD45 BV510 (HI30, BD Biosciences), anti-mouse CD43 BV605 (S7, BD Biosciences), anti-mouse CD93 BV650 (AA4.1, BD Biosciences), anti-human CD3 BV711 (UCHT1, BD Biosciences), anti-human CD19 BV786 (2H7, BD Biosciences), anti-human CD19 FITC (HIB19, BioLegend), anti-mouse B220/CD45R PerCP (RA3-6B2, BD Biosciences), anti-human CD10 PerCP-Cy5.5 (HI10a, BD Biosciences), anti-mouse CD4 BV711 (RM4-5, BioLegend), anti-mouse CD4 BV421 (GK1.5, BioLegend), anti-mouse CD103 APC & BV510 (2E7, BioLegend), anti-mouse Thy1.2 PerCP (53-2.1, BioLegend), anti-mouse CD197 PE (4B12, BioLegend), anti-mouse CD8a BV605 (53-6.7, BioLegend), and anti-mouse PD-1 PE-Cy7 (29F.1, BioLegend).

### Figure Generation

Figures were generated using BioRender, FlowJo 10 (Becton Dickinson & Company), Prism 9 (Dotmatics) and Illustrator 2021 (Adobe).

## AUTHOR CONTRIBUTIONS

K. S. Fitzpatrick produced antigen tetramers, designed, conducted, and analyzed murine cell and bead experiments; H. N. Degefu produced peptide:MHC tetramers, designed, conducted and analyzed peptide:MHC tetramer cell and bead experiments, generated figures and helped write portions of the manuscript; K. Poljakov conducted murine cell and bead experiments; M. G. Bibby conducted and analyzed BLI experiments; A. J. Remington produced tetramers, designed and conducted bead experiments; T.G. Searles produced peptide:MHC tetramers; M. D. Gray produced, purified, and validated antigens and antibodies; J. Boonyaratanakornkit conducted human cell sorting experiments to clone SARS-CoV-2-specific antibodies, conducted and analyzed BLI experiments, and edited the manuscript; P.C. Rosato designed peptide:MHC tetramer experiments, and wrote and edited portions of the manuscript; J.J. Taylor designed and analyzed experiments, generated figures, and wrote the manuscript.

## ACKNOWLEDGMENTS

We thank L. Stamatatos for CV30; M. Pepper and J. Netland for 204, 208, 211 and 215; M. J. McElrath for PBMCs from Seattle Area Control cohort; D. Koelle and A. Wald for COVID-19 PBMC samples from NCT04338360 and NCT04344977; B. Graham for SARS-CoV-2 S2P plasmid; Fred Hutch Flow Cytometry and Comparative Medicine Shared Resource staff for technical assistance; M. Lopez-Bernal, L. Yates, R. Putnam, and M. Gurtovnik for administrative assistance; S. Voght for manuscript editing; and D. Baumjohann, A. Broggi, K. Bruton, C. Castellanos, L. Chopp, C. Coelho, D. Glass, C. Hopp, J. Koenig, S. Langel, N. Mohamed, K. O’Connor, G. Rizzuto, C. Sundling, L. Swadling, P. Wilson, F. Wimmers, R. Yefet, and T. Yewdell for suggesting important citations on Twitter or through email; the bioMT Molecular Tools Core at Dartmouth for help with peptide:MHC monomer production; E. Ferris and M. Cole for use of their FPLC; the NIH tetramer core for *E.coli* expressing HLA-A2 heavy chain; D. Masopust for *E.coli* expressing mouse H2K^b^ and human β2m; and E. Newell for helpful discussions regarding MHC tetramer production. Research reported in this publication was supported by the National Institutes of Health under award numbers R01AI122912 to J.J. Taylor, T32GM095421 to K. Fitzpatrick, P20GM13132-07 to P.C. Rosato, Fast Grants award to J. Boonyaratanakornkit, and a Fred Hutch Evergreen award to J. Boonyaratanakornkit & J. J. Taylor.

